# Optimization of reference population for imputation of low-density SNPs panel for genomic prediction in Atlantic salmon

**DOI:** 10.1101/2025.11.17.688860

**Authors:** Clémence Fraslin, Serap Gonen, Solomon Boison, Matthew Baranski, Ashie Norris, Diego Robledo

## Abstract

In recent years many advances have been made towards developing cost-efficient low-density genomic tools for a wider implementation of genomic selection in aquaculture breeding programmes. Genotype imputation from very low-density (LD) SNP panels of just a few hundred markers to high-density (HD) SNP panels has become a promising strategy to reduce the cost of genotyping while maintaining accurate genomic prediction. The objective of this study is to assess the impact of the makeup of HD-genotyped reference populations on i) the accuracy of imputation for LD-genotyped individuals and ii) the accuracy of genomic prediction for three traits of importance in Atlantic salmon production: growth, resistance to cardiomyopathy syndrome and resistance to pancreas disease. An Atlantic salmon population genotyped with a 47K SNP array was used for the study, along with an *in silico* LD panel of 554 SNPs. Five reference population scenarios for imputation were tested, which could include only the parents of the candidates for selection, a combination of parents and candidates, or just candidates. All scenarios resulted in highly accurate imputation rates (over 80%) except when the HD reference population was only composed of selection candidates. Nonetheless, the accuracy of imputation barely had an impact on the accuracy of genomic prediction, as the imputed datasets performed very similarly to the HD-panel. Adding a proportion of the offspring to the reference population, in addition to the parents, did not result in any benefit in terms of genomic prediction. Imputation is a cost-effective and robust option for genomic selection in aquaculture.

## Introduction

In modern selective breeding programmes, genomic selection is used to identify the best candidates based on their genomic breeding values that are predicted from their own genotypes and from genotype and phenotype information obtained from a related training population (Goddard and Hayes, 2007; Meuwissen, 2009). Compared to a pedigree-based approach, for traits that cannot be measured in the selection candidates, this method allows to utilise within family genetic variance instead of relying on family overall performance. Genomic selection is increasingly applied in aquaculture and has been shown to be more accurate than pedigree-based approaches for most traits under selection, especially when measured on close relatives of the candidates (sib-selection) as reviewed recently by (Houston et al., 2020; Robinson et al., 2022; You et al., 2020). Increasing the accuracy of prediction is a key factor for selective breeding as this is directly correlated to a higher genetic gain (Δ*G*) according to the breeder’s equation for genetic gain: 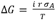 where r is the prediction accuracy, i the selection intensity, *σ*_*A*_ the additive genetic variance and T the generation interval.

Aquaculture breeding programs rely on a high number of full-sib families (∼200) with large number of offspring, which has facilitated the uptake of genomic selection due to the possibility of having a close relationship between selection candidates and the training population. The training population(s) is usually composed of full and half-sibs (Fraslin et al., 2022), which can be routinely phenotyped in diverse environments, including farms settings. While these sib-testing schemes tend to result in high prediction accuracies, a major bottleneck for a wide implementation of genomic selection in aquaculture is the need to genotype a large training population to obtain accurate predictions and a large number of candidates to achieve sufficient selection intensity. Considering current genotyping prices, this can be prohibitive for many farmers (Boudry et al., 2021). Decreasing the cost of individual genotyping would allow a wider implementation of genomic selection in aquaculture breeding programmes, making it affordable for small and medium-sized companies. For larger companies that have already been implementing genomic selection in their breeding programmes (Norris, 2017; Zenger et al., 2019), reducing the individual cost of genotyping would allow the genotyping of additional candidates, enabling an increase in selection intensity thus increasing genetic gain. In recent years, for various aquaculture species, there has been a major focus on designing optimised low-density (LD) panels with between 1,000 and 6,000 SNPs, which have produced breeding value estimates as accurate as those of medium or high-density panels for a large number of traits (Griot et al., 2021; Kriaridou et al., 2020; Peñaloza et al., 2022; Song et al., 2022; Song and Hu, 2022; Vallejo et al., 2021). Additionally, recent studies have shown that genotyping the selection and training populations at very low densities of a few hundred SNPs can be combined with genotype imputation using a reference population genotyped with medium-density (MD) or high-density (HD) panels to obtain genomic predictions as accurate as those obtained with all fish genotyped at MD/HD (Dufflocq et al., 2019; Fraslin et al., 2023; Griot et al., 2021; Kriaridou et al., 2023; Mastrochirico-Filho et al., 2024; Tsairidou et al., 2020; Yoshida et al., 2019). Genotype imputation relies on linkage disequilibrium between SNPs in close proximity within a chromosome, which are likely to be inherited together. Close relatives (parents-offspring or full-sibs) will share long haplotypes, while distant relatives will share shorter haplotypes that will have been broken down through recombination (Kong et al., 2008). Within a breeding programme, the reference population for imputation is key, however its makeup has not been investigated in an aquaculture context, both in terms of number of individuals and their relationship to the imputed individuals. Indeed, most of the recently published studies investigating the optimization of LD-panels performed imputation using the HD-genotyped parents as reference population for imputation.

In this study we compared different scenarios to optimize the construction of the reference population for an accurate imputation of LD-genotypes in an Atlantic salmon (*Salmo salar*) commercial population, and we investigated the accuracy of prediction using imputed data for three traits of interest, resistance to cardiomyopathy syndrome, resistance to pancreas disease and growth.

## Material and methods

### Phenotypes and genotypes

Atlantic salmon phenotypes, genotypes and pedigree data were obtained from MOWI Norway’s 2019 year-class (YC), composed of 336 full-sib families (some of them half-sibs) from 134 sires (from YC2012 to YC2015) and 221 dams (from YC2015 to YC2016). The full-sib families were phenotyped for three traits: resistance to cardiomyopathy syndrome (CMS, n=1,172), resistance to pancreas disease (PD, n=1,500) and growth (n=3,108) measured as gutted weight (GW) at harvest.

The disease challenges to measure resistance to CMS and resistance to PD were performed as follows.

For the PD challenge, about 4,500 fry were transported with water and oxygen to VESO Vikan (Norway) in May 2018. After approximately 3 weeks of acclimatisation the fish were challenged with PD virus (SAV2) by a modified cohabitant model as described in (Gonen et al., 2015). Tissue samples of dead fry were sampled daily, and the challenge was ended after 82 days when mortality had levelled off (the final mortality was 32%). For the purposes of this study the genotypes and phenotypes of 1500 randomly selected fish (equal numbers survivors and mortalities) were made available. Resistance to PD (strain SAV2) was measured as binary survival (1 dead, 2 alive).

For the CMS challenge, about 1,700 smolts were transported to VESO Vikan (Norway) in May 2019. All surviving fish were intraperitoneally injected with PMC virus (Boison et al., 2019). At the end of the challenge period that lasted approximately 8 weeks, hearts were removed from all fish and sent for histopathological examination and scoring. Only the phenotypes of fish surviving past a defined period were included in the current analysis (1,172). Resistance was measured as a histology score of the atrium (heart) tissue, ranging from 0 (best score, absence of histopathological findings) to 4 (worst score, lesions in more than 75% of the tissue and moderate to severe leukocyte infiltration).

All the YC2019 fish and most of the parents (21 parents were missing) were genotyped with the ThermoFisher Axiom 57K SNP array (NOFSAL03 containing 55,735 SNPs, Mowi-owned data) developed by Nofima in collaboration with Mowi and SalmoBreed (now Benchmark Genetics). After standard quality controls (minor allele frequency ≥ 0.05, SNP and individual call rate ≥ 95% and departure from Hardy-Weinberg equilibrium p ≤ 1.10^-15^), 5,873 fish genotyped for 47,061 SNPs remained; 1,094 of the fish had phenotypes for resistance to CMS, 1,448 for resistance to PD 2,998 for growth, and 334 parents were kept. A low-density (LD) panel was created *in silico* by masking the genotypes of YC2019 fish to only keep the information of 554 SNPs from a pre-established list with SNPs carefully selected by Mowi (Dagnachew, personal communication).

### Imputation and optimization of reference population

The “missing” genotypes (masked *in silico*) of individuals from the YC2019, considered as the LD-genotyped target population, were imputed from the LD-panel (554 SNPs) to the high density (HD, 47K SNPs) using a combined pedigree and population based approach implemented in the FImpute software (v.3, (Sargolzaei et al., 2014)). The optimal design of the HD-genotyped reference population was tested in five different scenarios presented below and in Table 1:

**Table 1.**
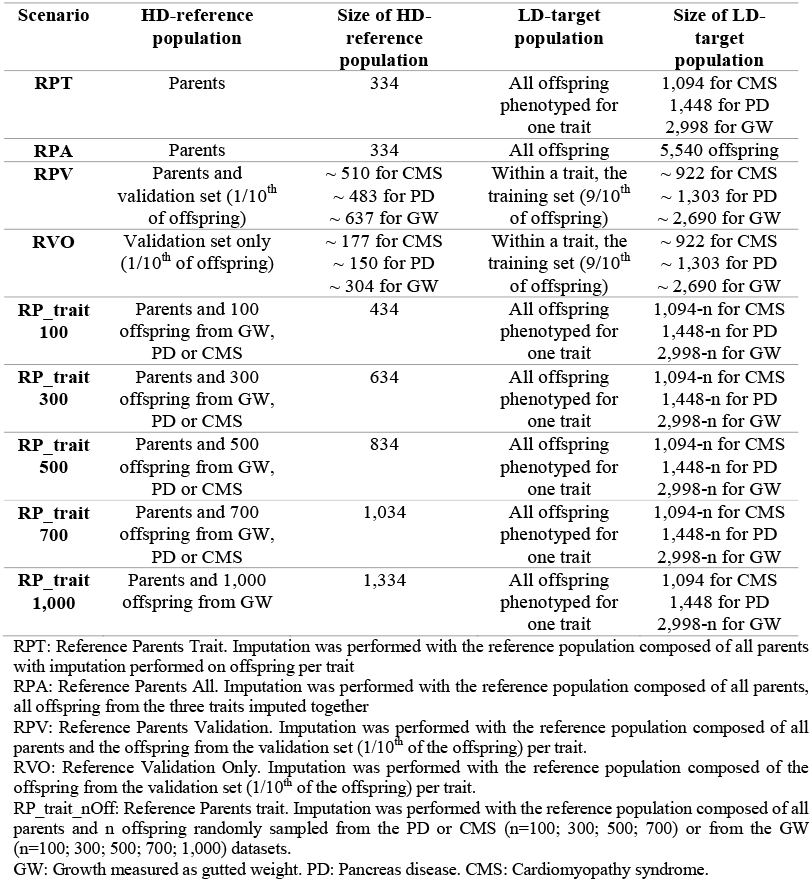
Different scenarios for imputation of the LD-genotyped target population.

| Scenario | HD-reference population | Size of HD-reference population | LD-target population | Size of LD-target population |
| --- | --- | --- | --- | --- |
| **RPT** | Parents | 334 | All offspring phenotyped for one trait | 1,094 for CMS<br>1,448 for PD<br>2,998 for GW |
| **RPA** | Parents | 334 | All offspring | 5,540 offspring |
| **RPV** | Parents and validation set (1/10<sup>th</sup> of offspring) | ~ 510 for CMS<br>~ 483 for PD<br>~ 637 for GW | Within a trait, the training set (9/10<sup>th</sup> of offspring) | ~ 922 for CMS<br>~ 1,303 for PD<br>~ 2,690 for GW |
| **RVO** | Validation set only (1/10<sup>th</sup> of offspring) | ~ 177 for CMS<br>~ 150 for PD<br>~ 304 for GW | Within a trait, the training set (9/10<sup>th</sup> of offspring) | ~ 922 for CMS<br>~ 1,303 for PD<br>~ 2,690 for GW |
| **RP_trait 100** | Parents and 100 offspring from GW, PD or CMS | 434 | All offspring phenotyped for one trait | 1,094-n for CMS<br>1,448-n for PD<br>2,998-n for GW |
| **RP_trait 300** | Parents and 300 offspring from GW, PD or CMS | 634 | All offspring phenotyped for one trait | 1,094-n for CMS<br>1,448-n for PD<br>2,998-n for GW |
| **RP_trait 500** | Parents and 500 offspring from GW, PD or CMS | 834 | All offspring phenotyped for one trait | 1,094-n for CMS<br>1,448-n for PD<br>2,998-n for GW |
| **RP_trait 700** | Parents and 700 offspring from GW, PD or CMS | 1,034 | All offspring phenotyped for one trait | 1,094-n for CMS<br>1,448-n for PD<br>2,998-n for GW |
| **RP_trait 1,000** | Parents and 1,000 offspring from GW | 1,334 | All offspring phenotyped for one trait | 1,094 for CMS<br>1,448 for PD<br>2,998-n for GW |

RPT: Reference Parents Trait. Imputation was performed with the reference population composed of all parents with imputation performed on offspring per trait
RPA: Reference Parents All. Imputation was performed with the reference population composed of all parents, all offspring from the three traits imputed together
RPV: Reference Parents Validation. Imputation was performed with the reference population composed of all parents and the offspring from the validation set (1/10<sup>th</sup> of the offspring) per trait.
RVO: Reference Validation Only. Imputation was performed with the reference population composed of the offspring from the validation set (1/10<sup>th</sup> of the offspring) per trait.
RP\_trait\_nOff: Reference Parents trait. Imputation was performed with the reference population composed of all parents and n offspring randomly sampled from the PD or CMS (n=100; 300; 500; 700) or from the GW (n=100; 300; 500; 700; 1,000) datasets.
GW: Growth measured as gutted weight. PD: Pancreas disease. CMS: Cardiomyopathy syndrome.

1. RPT (Reference Parents per Trait) where the reference population was composed of only the 334 parents. Three independent target populations were created (one for each trait) each of them containing all offspring with phenotypes recorded for one trait. All three target populations were imputed independently (i.e., imputation was run three times: one for GW, one for PD and one for CMS). This scenario corresponds to a scenario where the breeder can perform genotype imputation and estimate breeding values through genomic prediction every time a new phenotype is available without having to wait for a complete dataset of the population with all phenotypes.
2. RPA (Reference Parents All) where the reference population was also composed of only the 334 parents. The target population was composed of all 5,540 phenotyped offspring, imputed to HD in a single run. This scenario was created to see if the size of the target population had an impact on imputation. This scenario corresponds to a scenario where the breeder needs to wait until all phenotypes are available to perform genotype imputation and genomic prediction on the whole population simultaneously.
3. RPV (Reference Parents Validation) where the reference population was composed of all the parents and 1/10^th^ of the offspring within one trait, corresponding to the fish used as validation set for the genomic prediction analysis. All traits were imputed separately and the target population was composed of 9/10^th^ of the fish with a phenotype recorded for a given trait, corresponding to the training population for the genomic prediction analysis. This scenario corresponds to a scenario where the breeder wants to genotype the selection candidates (validation population) with an HD-panel.
4. RVO (Reference Validation Only) where the reference population was composed of 1/10^th^ of the offspring within one trait, corresponding to the fish in the validation set for the genomic prediction analysis. All traits were imputed separately and the target population was composed of 9/10^th^ of the fish with a phenotype recorded for a given trait, corresponding to the training population for the genomic prediction analysis. This scenario corresponds to a scenario where the breeder wants to genotype the selection candidates (validation population) with an HD-panel and there are no genotypes available for the parental generation.

Finally, it was decided to test a fifth scenario that better reflects industry practices. If part of the offspring would have to be genotyped with the HD-panel and included in the reference population with the HD-genotyped parents, the fish phenotyped for a disease resistance trait are the most likely to be included in the reference as such phenotypes are frequently obtained on early life stages. However, we decided to test the scenario for all traits. Thus, three new scenarios, RP_GW_n, RP_PD_n and RP_CMS_n (RP_trait_nOff = Reference Parents trait n offspring) were designed in which the reference population for imputation was composed of all the parents and different number (n) of the fish phenotyped (n=100; 300; 500; 700 for PD and CMS and up to n=1,000 for GW). The n phenotyped fish to be included in the reference population were randomly sampled and 10 replicates were created for each n to minimise bias due to sampling.

The quality of imputation was estimated for each scenario using two values. First, the accuracy of imputation was estimated, for each individual, as the Pearson’s correlation between the true genotype and the imputed genotype. Then, the proportion of correctly imputed genotypes was estimated, for each SNP, as the number of correctly imputed genotypes divided by the number of imputed genotypes. In both cases, the genotypes were coded as allele count (0,1,2), the SNPs that were on the LD panel (*i*.*e*., not imputed) and the SNPs with a missing genotype in the HD-panel (*i*.*e*., SNPs for which we did not know the true genotype) were not used to assess the quality of imputation. Thus, the number of imputed SNPs was different from the number of SNPs in the HD-panel (47,061) and was unique for each marker on the SNP array and each individual.

### Genetic parameters and breeding values estimation

Variance components and estimated breeding values (EBV) of individuals were computed for each trait using BLUPF90 software (Misztal et al., 2002) following the mixed linear BLUP-animal model:

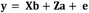

Where the vector of phenotype (**y**) was explained by a vector of fixed effect (**b**), a vector of random additive genetic effect (**u**) and a vector of residual effect (**e**). **X** and **Z** are the incidences matrices for fixed effect and animal genetic effect, respectively. The vector of residual effect (**e**) follows a normal distribution 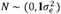 with 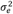 the residual variance and **I** the identity matrix. The additive genetic effect (***a***) follows a normal distribution 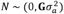 with 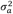 the additive genetic variance and **G** the relationship matrix as described in (VanRaden, 2008). For the variance component analysis, the relationship matrix used in the model was the genomic relationship matrix computed with the genotype information from all markers in the HD-panel. Estimated breeding values (EBV) and Genomic estimated breeding values (GEBV) of individuals were computed using pedigree and genomic BLUP (PBLUP and GBLUP) with i) only pedigree information for the PBLUP and genotypes from the ii) HD-panel, iii) LD-panels and iv) imputed LD-panels. The only significant fixed effect that was included in the model was the sex of the fish for the growth phenotype (GW). No fixed effect or covariates were included in the model for the two disease resistance traits.

Variance components were estimated using the airemlf90 package and (G)EBVs were estimated using the blupf90+ package (Misztal et al., 2002).

### Accuracy of genomic prediction

The accuracy of genomic prediction for all three traits was estimated using Monte-Carlo “leave-one-group-out” method by removing the known phenotype from 1/10^th^ of the fish to create a validation group, and then using the information from the remaining 9/10^th^ fish in the training group to predict the (G)EBVs of the validation group using the model described in the section above. This procedure was repeated 10 times. Accuracy of prediction was computed as the mean over the 10 replicates of the Pearson’s correlation coefficient between the (G)EBV and the true phenotypes of fish in the validation group, divided by the square root of the genomic based heritability estimated with the HD-panel (Legarra et al., 2008). The inflation coefficient was derived as the regression coefficient of the true phenotypes on the (G)EBVs. This coefficient is expected to be equal to 1 in the absence of selection bias, while a value below 1 represents inflation (over-dispersion of EBVs) and a value above 1 deflation (under-dispersion of EBVs). The significance of the differences between imputation accuracies and prediction accuracies obtained with different panel densities and before or after imputation was tested using pairwise Dunn tests with a Benjamini-Hochberg correction for multiple comparison using the dunn.test R package (v 1.3.5).

## Results

### Phenotypes and genetic parameters estimate

After genotype quality control, 1,094 of the fish had phenotypes for resistance to CMS, and the scores were distributed as 0 n=163, 1 n=418, 2 n=395, 3 n=107, 4 n=11. 1,488 individuals remained for resistance to PD, with a survival rate after infection of 49% (by design as genotyped fish were selected to contain 50% survivors and 50% dead). Finally, 2,998 individual phenotypes for growth were retained, measured as gutted weight (GW; mean = 4.87 kg ± 0.833 sd).

Heritability estimates for each trait using the HD-panel along with variance component estimates are presented in table 2.

**Table 2.** Estimates of genomic parameters for resistance to pancreas disease, to cardiomyopathy syndrome and gutted weight.

| | $h^2 \pm se$ | $\sigma_a^2 \pm se$ | $\sigma_p^2 \pm se$ | $\sigma_e^2 \pm se$ |
| --- | --- | --- | --- | --- |
| **GW** | 0.43 ± 0.031 | 0.29 ± 0.029 | 0.68 ± 0.026 | 0.39 ± 0.014 |
| **PD** | 0.46 ± 0.043 | 0.12 ± 0.015 | 0.26 ± 0.012 | 0.14 ± 0.008 |
| **CMS** | 0.47 ± 0.049 | 0.38 ± 0.052 | 0.81 ± 0.041 | 0.43 ± 0.033 |

h<sup>2</sup>: genomic heritability calculated as $\sigma_a^2 / (\sigma_a^2 + \sigma_e^2)$ . $\sigma_a^2$ : Additive genetic variance. $\sigma_p^2$ : Phenotypic variance calculated as $(\sigma_a^2 + \sigma_e^2)$ . $\sigma_e^2$ : Residual variance.

### Accuracy of imputation

The accuracy of imputation was overall very high with an average accuracy over 0.8 for 4 out of 5 scenarios, a value that is usually considered as the threshold for an accurate imputation; results for the first four scenarios are presented in Figure 1 and results for the RP_trait_nOff scenario are presented in Figure 2. For all traits, the scenario with the HD-genotyped reference population composed of only the offspring from the validation set (RVO scenario) resulted in a significantly lower imputation accuracy than all the other scenarios, with average accuracies of 0.79 (± 0.05 sd), 0.75 (± 0.06 sd) and 0.76 (± 0.06 sd) for GW, PD and CMS, respectively. The proportion of correctly imputed SNP was also significantly lower for this RVO scenario compared to other scenarios, with average proportions of 84.5% (± 3.4 sd), 81.8% (± 4.0 sd) and 82.0% (± 4.0 sd) for GW, PD and CMS, respectively. The highest accuracy and proportion of correctly imputed SNP were obtained when the HD-genotyped reference population was composed of the parents and the validation set (RPV scenario). Average accuracies per individual and proportion of correctly imputed SNPs were 0.93 (± 0.07 sd) and 94.4% (± 5.3 sd) for GW, 0.91 (± 0.08 sd) and 93.0% (± 5.9 sd) for PD and 0.92 (± 0.07 sd) and 93.3% (± 5.1 sd) for CMS. For all three traits the two scenarios with only the parents in the HD-genotyped reference population (RPT and RPA) performed similarly, with an average accuracy for imputation ranging between 0.87 and 0.88, while the proportion of correctly imputed SNPs ranged between 89.5% and 90.5% so the size of LD-target population – or at least adding more animals of the same families - had very limited impact on the imputation accuracy (*p-values* < 5.0e-4).

**Figure 1.**
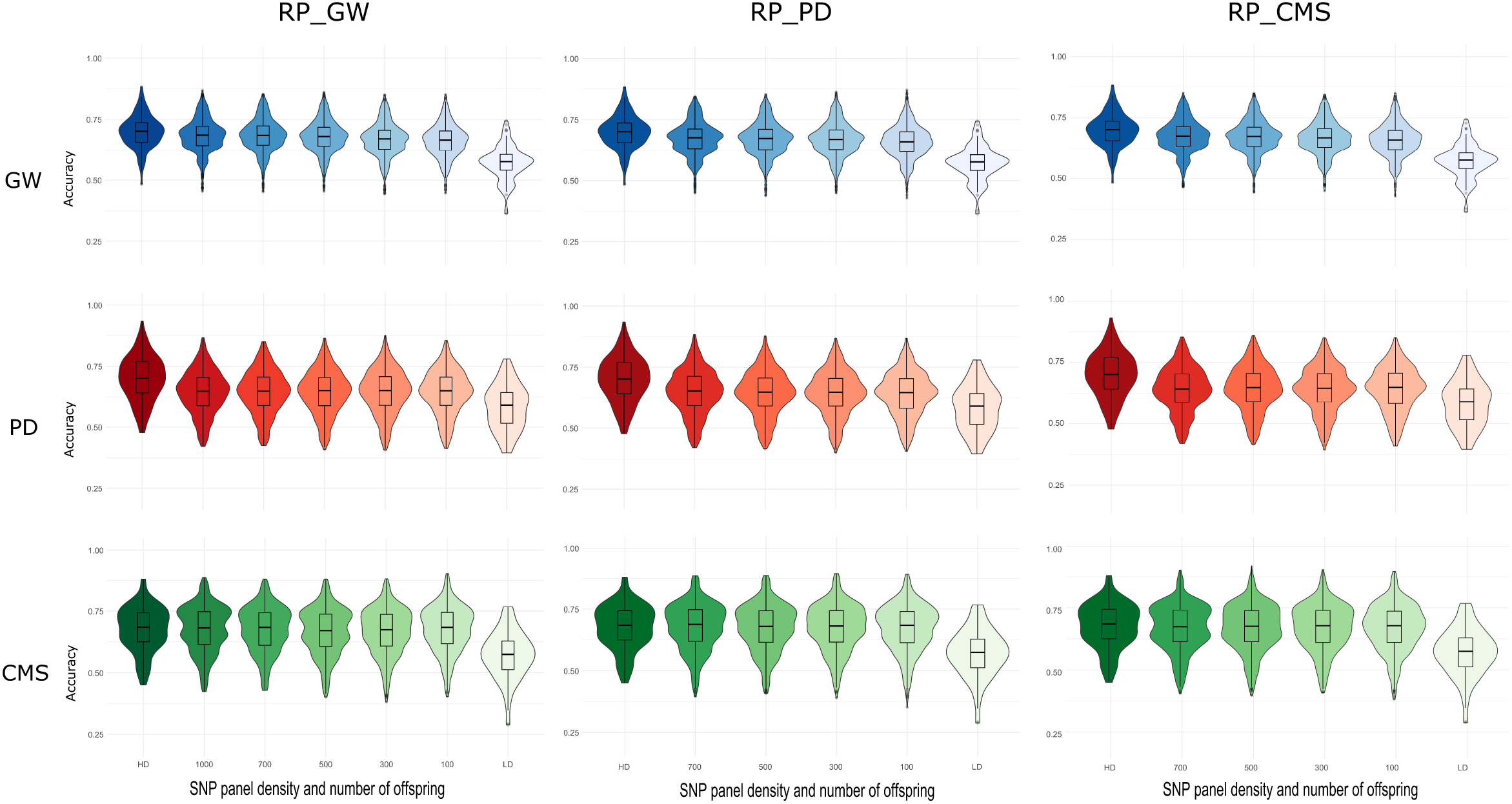
Accuracy of imputation per individual for three traits when 554 SNPs are imputed to 47K SNPs under four imputation scenarios. Accuracy measured as the Pearson correlation between true and imputed genotypes (coded 0,1 or 2). RPT: Reference Parents Trait. Imputation was performed with the reference population composed of all parents with imputation performed on offspring per trait. RPA: Reference Parents All. Imputation was performed with the reference population composed of all parents, all offspring from the three traits imputed together. RPV: Reference Parents Validation. Imputation was performed with the reference population composed of all parents and the offspring from the validation set (1/10^th^ of the offspring) per trait. RVO: Reference Validation Only. Imputation was performed with the reference population composed of the offspring from the validation set (1/10^th^ of the offspring) per trait. GW: Growth measured as gutted weight. PD: Pancreas disease. CMS: Cardiomyopathy syndrome.

**Figure 2.**
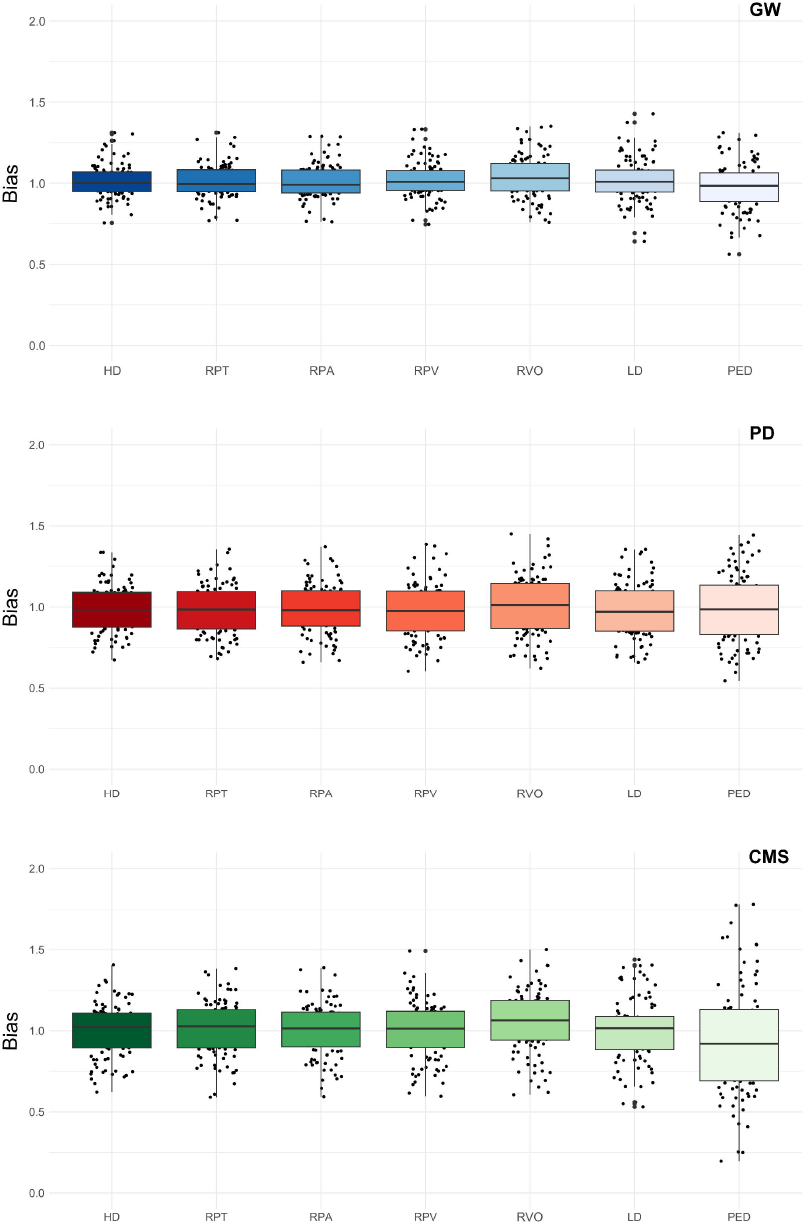
Accuracy of imputation per individual for three traits when 554 SNPs are imputed to 47K SNPs when different number of offspring from the GW, PD or CMS datasets are added to the HD-genotyped reference population for imputation. Accuracy measured as the Pearson correlation between true and imputed genotypes coded 0,1 or 2 RP_trait_nOff: Reference Parents trait. Imputation was performed with the reference population composed of all parents and n (100 to 700 or 1000) offspring randomly sampled from the GW, PD or CMS datasets. GW: Growth measured as gutted weight. PD: Pancreas disease. CMS: Cardiomyopathy syndrome.

For the scenario where a random proportion of fish from the growth dataset were added to the reference panel (RP_trait_nOff with n=100 to n=700 or 1,000 offspring), the quality of imputation, measured as both correlation or proportion of correctly imputed genotypes, increased slightly with the number of individuals in the reference population for all three traits (Figure 2). For all three traits, the lowest number of correctly imputed genotypes per individual were obtained when only 100 offspring were added to the reference panel. In this scenario, regardless of the origin of those offspring (RP_GW, RP_CMS, RP_PD), the imputations accuracy and the proportion of correctly imputed SNPs ranged between 0.90 and 92.0% (for GW imputed in RP_CMS) to 0.91 and 92.7% (for CMS imputed in RP_GW), respectively. The highest proportion of correctly imputed genotypes was always obtained with the largest reference population i.e., with 1000 offspring from the GW dataset (RP_GW_1000), independently of the trait of the population to impute, and with 700 offspring from the same phenotype as the population to impute. In detail, for individuals of the GW phenotype dataset the highest proportion of correctly imputed genotypes was obtained with the RP_GW_1000 and RP_GW_700 scenarios (95.8% and 95.5%, respectively; *p-value* = 0.032). For the CMS dataset, the highest proportion of correctly imputed genotypes (95.2%) was obtained with the RP_GW_1000 and RP_CMS_700 scenarios (non-significant differences, see supplementary Table S1). Finally, for the PD dataset, the highest proportion of correctly imputed genotypes was obtained with the RP_GW_1000 and RP_PD_700 (95.43% and 95,37% respectively; *p-value* < 0.001).

Imputation is considered successful with an accuracy over 0.8, while if the accuracy drops below 0.5 it is usually considered that the imputation is not better than randomly assigning genotypes. The number and proportion of individual with poor (accuracy below 0.8) and inaccurate (below 0.5) imputation are presented in supplementary Table S2. Interestingly, when only the validation group was used as HD-genotyped reference population (RVO), most of the fish phenotyped for PD (94%) and CMS (89%) and about half of the fish from GW (50%) had a poor accuracy of imputation (below 0.8), however only one individual phenotyped for GW had an imputation accuracy lower than 0.5.

The RPT, RPA and RP_trait_100 scenarios performed similarly in terms of proportion of accurately imputed individuals (with an accuracy over 0.8) with around 88%, 87% and 90% of the individual correctly imputed for GW, PD and CMS, respectively. Within those scenarios, almost all the poorly imputed animals had one or both parents missing from the HD-reference panel and their individual imputation accuracy was improved when siblings were added to the HD-reference panel (RVO, RPV and all RP_trait_nOff scenarios). The scenario that yielded the highest proportion of accurately imputed individuals was RP_PD_700 with 94.7%, 94.8% and 95.8% of individuals that were accurately imputed for GW, PD and CMS, respectively.

### Accuracy of genomic prediction

For all three traits, genomic predictions were always more accurate than pedigree-based predictions and the accuracy of predicted GEBVs increased significantly with imputation compared to the un-imputed LD-panel (Figure 3, supplementary Table S3). The two scenarios PRA and PRT, which only differed in whether the full dataset was imputed at once (RPA) or per trait (RPT) performed the same for all three traits (non-significant differences, see supplementary Table S3).

**Figure 3.**
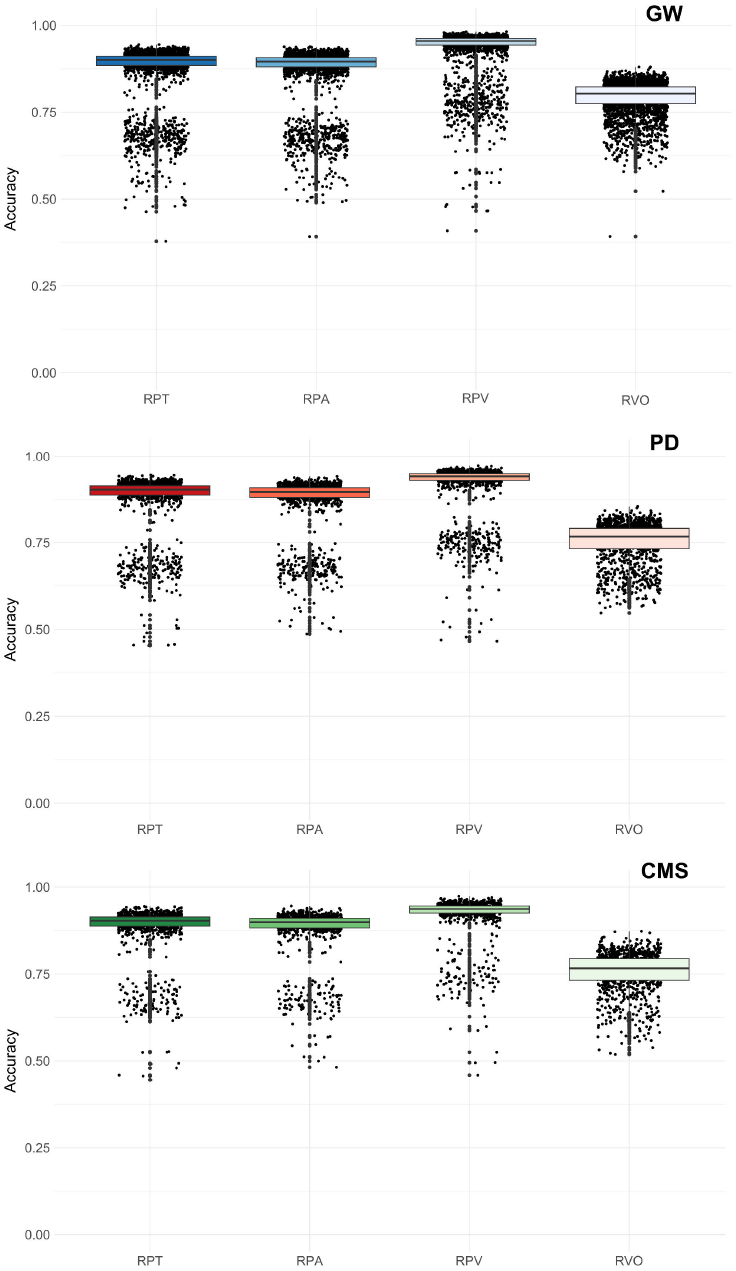
Accuracy of genomic predictions for three traits in an Atlantic salmon population for various density panels and imputation scenarios. HD: Prediction obtained with the high-density panel (47K SNPs). RPT: Reference Parents Trait. Imputation was performed with the reference population composed of all parents with imputation performed on offspring per trait. RPA: Reference Parents All. Imputation was performed with the reference population composed of all parents, all offspring from the three traits imputed together. RPV: Reference Parents Validation. Imputation was performed with the reference population composed of all parents and the offspring from the validation set (1/10^th^ of the offspring) per trait. RVO: Reference Validation Only. Imputation was performed with the reference population composed of the offspring from the validation set (1/10^th^ of the offspring) per trait. LD: Prediction obtained with the un-imputed low-density panel (554 SNPs). PED: Prediction obtained with a pedigree-based approach. GW: Growth measured as gutted weight. PD: Pancreas disease. CMS: Cardiomyopathy syndrome.

The comparison between the accuracy of prediction obtained from the HD panel and from the imputed panel varied slightly across the traits. For GW, the HD-panel performed significantly better than all imputed scenario except when the HD-genotyped reference population for imputation included the parents and offspring from the validation set (RPV), which was nearly as good as the HD-panel (*p*-value = 0.053) to predict GEBV. For this trait, the accuracy obtained after imputation with RPV was similar to those obtained with the scenarios RPA and RPT (*p-value* > 0.1). For resistance to PD, the predictions obtained with the HD panel were always significantly more accurate than the predictions obtained with imputed panels (supplementary Table S3), no matter which imputation scenario was employed. All imputation scenarios resulted in similar accuracy of prediction, and all performed better than the LD-panel and the pedigree-based predictions. For resistance to CMS, all imputation scenarios were as good as the HD-panel and as good as each other to predict the GEBV of the fish (differences not significant). Imputation significantly increased the accuracy of prediction compared to genomic prediction obtained with the LD-panel and to pedigree-based predictions.

Imputation did not significantly inflate or deflate the GEBVs compared to the GEBVs obtained with the HD-panel (*p*-values > 0.2). For all imputation scenarios, except the pedigree-based accuracy for resistance to CMS, there was on average little bias (Figure 4). For all traits, the pedigree-based prediction was slightly more biased than the genomic based predictions no matter the SNP density (not significant) with in average a regression coefficient value below 1 indicating an over-dispersion (inflation) of the EBVs. For all traits, GEBVs obtained after imputation with the RVO scenario were on average slightly under-dispersed.

**Figure 4.**
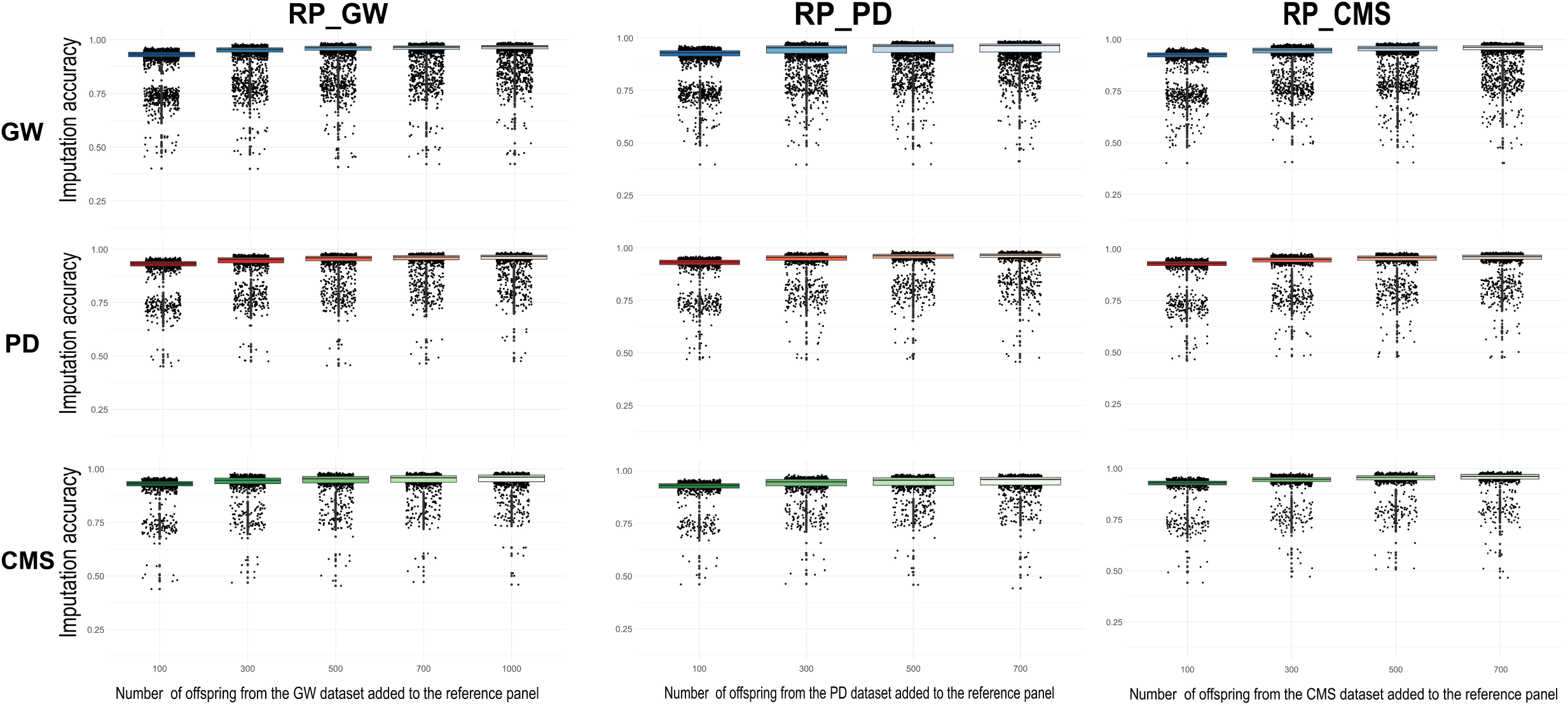
Bias of genomic predictions for three traits in an Atlantic salmon population for various density panels and imputation scenarios. HD: Prediction obtained with the high-density panel (47K SNPs). RPT: Reference Parents Trait. Imputation was performed with the reference population composed of all parents with imputation performed on offspring per trait. RPA: Reference Parents All. Imputation was performed with the reference population composed of all parents, all offspring from the three traits imputed together. RPV: Reference Parents Validation. Imputation was performed with the reference population composed of all parents and the offspring from the validation set (1/10^th^ of the offspring) per trait. RVO: Reference Validation Only. Imputation was performed with the reference population composed of the offspring from the validation set (1/10^th^ of the offspring) per trait. LD: Prediction obtained with the un-imputed low-density panel (554 SNPs). PED: Prediction obtained with a pedigree-based approach. GW: Growth measured as gutted weight. PD: Pancreas disease. CMS: Cardiomyopathy syndrome.

Figure 5 shows the accuracy of genomic prediction when 100 to 700 or 1,000 random offspring from the different datasets are added to the reference population for imputation. For all traits the accuracy of genomic prediction was always improved in the RP_trait_nOff scenario compared to the one obtained with the LD panel, it was similar to the RPT scenario with only the parents in the reference population and was slightly lower than the one obtained with the HD panel. For GW, the accuracy increased as more offspring from the same dataset were added to the reference population. The maximum accuracy of genomic prediction was reached with 700 and 1,000 offspring added, but remained lower than the accuracy obtained with the HD-panel (although the difference was not significant, see supplementary Table S4a). For both disease resistance traits there was no significant improvement of the accuracy of genomic prediction when 100 or 700/1,000 fish from any dataset were added to the reference population for imputation (Figure 5 and supplementary Tables S4b and S4c).

**Figure 5.**
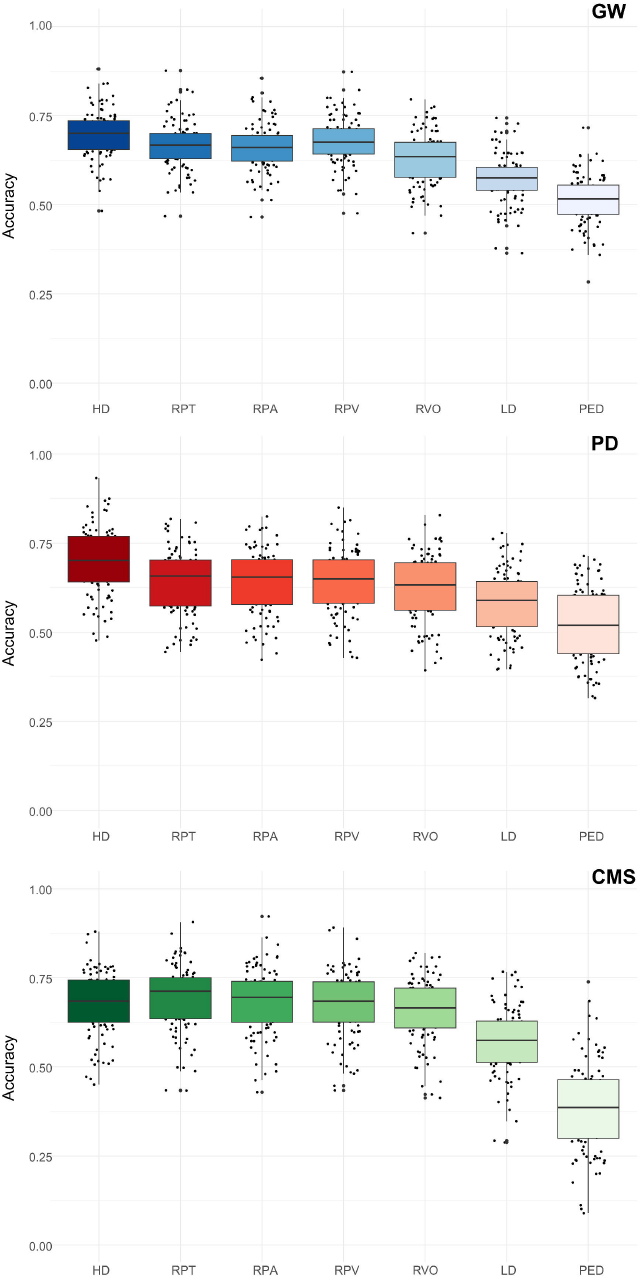
Accuracy of prediction for three traits when different number of offspring from the different datasets were added to the reference population for imputation. In column A: Reference Parents GW. Imputation was performed with the reference population composed of all parents and n (100, 300, 500, 700 or 1000) offspring randomly sampled from the growth dataset. In column B: Reference Parents PD. Imputation was performed with the reference population composed of all parents and n (100, 300, 500 or 700) offspring randomly sampled from the PD dataset. In column C: Reference Parents CMS. Imputation was performed with the reference population composed of all parents and n (100, 300, 500 or 700) offspring randomly sampled from the CMS dataset. HD: Prediction obtained with the high-density panel (47K SNPs). RPT: Reference Parents per trait. Imputation was performed per trait with the reference population composed of all parents and no offspring. LD: Prediction obtained with the un-imputed low-density panel (554 SNPs). GW: Growth measured as gutted weight (row 1, in blue). PD: Pancreas disease (row 2, in red). CMS: Cardiomyopathy syndrome (row 3, in green).

In particular, for resistance to PD, all accuracies obtained were similar to those of the other imputed panels, with significantly lower prediction accuracy than that of the HD-panel (see supplementary Table S4b). For resistance to CMS, the accuracy varied a little depending on the number of offspring added to the reference population, but still remained similar to the accuracy obtained with the HD-panel (non-significant differences) and to that of the other imputation scenarios (see supplementary Table S4c).

## Discussion

Numerous recent studies have shown that combining low density genotyping and genotype imputation using a reference population genotyped at a higher density (usually tens of thousands of SNPs) is a powerful strategy to reduce the cost of genotyping in aquaculture breeding programmes. There is now an extensive body of literature on various fish species that investigated different SNP sampling methods to create in silico low density SNP arrays that are later imputed with different imputation software (Dufflocq et al., 2019; Fraslin et al., 2023; Griot et al., 2021; Kriaridou et al., 2023, 2020; Mastrochirico-Filho et al., 2024; Tsairidou et al., 2020; Vallejo et al., 2021; Yoshida et al., 2019). Most of these studies reached a consensus that the lowest optimal SNP density without imputation varies between 1,000 and 5,000 SNPs depending on the fish species, and that just a few hundred SNPs combined with imputation yield genomic predictions that are as accurate as the ones obtained with HD-SNP arrays.

In the current study we aimed to characterised the impact of realistic genotyping and imputation strategies on the accuracy of genomic prediction for three traits of major interest in Atlantic salmon farming. For instance, we investigated, *in silico*, the effect of the makeup of the HD-genotyped used as reference population for imputation on the performance of both imputation and genomic prediction. The low-density SNP panel was an in-house 500 SNP array from the salmon farming company Mowi. Five different scenarios were tested, two where only the parents were genotyped at HD-level and included in the reference population for imputation, two where the parents and a proportion of offspring were genotyped at HD, and one with only offspring in the HD-genotyped reference population. Imputation was very accurate in all scenarios with the parents included in the HD-reference population. Having only offspring from the validation set in the HD-reference population resulted in the lowest imputation accuracy for all three traits. We confirmed that genotype imputation significantly increased the accuracy of prediction compared to the 500 SNPs from the LD panel, reaching accuracies similar to the one obtained with the HD-panel (47K SNPs). The makeup of the reference population for imputation had little impact on the accuracy of genomic prediction.

### Accuracy of imputation

The accuracy of imputation, evaluated for each individual and each SNP, was overall very high, with an average individual accuracy over 0.8 and average proportion of correctly imputed SNPs above 90% in all scenarios. Considering the LD-panel is composed of only 554 SNPs, representing just 1.2% of the HD-panel (47,061 SNPs), imputation was remarkably successful. The highest accuracy of imputation was obtained when half and full-sibs of the LD-target population were added to the parents to form the HD-genotyped reference population (RPV and RP_trait_nOff scenarios). Imputation accuracy values obtained in the current study are within the range of previously reported accuracies for Atlantic salmon using similar size LD-panels and FImpute, independently of the size of the HD panel (Kriaridou et al., 2023; Tsairidou et al., 2020; Yoshida et al., 2018). This is expected as parents and offspring share long haplotype blocks at a low frequency, which increases the accuracy of imputation with FImpute as it uses overlapping sliding windows from longer haplotypes to shorter haplotypes (Sargolzaei et al., 2014). The family-based approach for imputation is highly accurate when close relatives of the target LD-individuals are used as HD-reference population no matter the species (Larmer et al., 2014; Sargolzaei et al., 2014; Yoshida et al., 2018; Zhang and Druet, 2010).

The lowest accuracy of imputation was obtained when imputation was performed in the RVO scenario that uses a HD-reference population composed only of half and full-sibs of the target population (i.e. no parents). Mainly, with this approach less individuals were imputed with a high accuracy (above 0.8; see Table S2), which might be explained by the less efficient phasing and identification of haplotypes due to the absence of fish from the parental generation in the HD-reference population. Indeed, genotype information from sires and dams is very important for phasing especially for very sparse LD-panels as family information is used to detect long-range haplotypes accurately (Sargolzaei et al., 2014). Interestingly, when a portion of the offspring were included in the HD-reference population (RPV and RPG_n) fewer individuals were poorly imputed (accuracy of imputation below 0.8) compared to scenarios with only HD-parents (see Table S2). The individuals with low or bad imputation accuracy in RPT and RPA scenarios usually had one or both parents missing (not present in the genotyping data); adding closely related siblings in the HD-reference population allowed to more accurately build the haplotypes for imputation increasing their imputation accuracy when the parents were not genotyped. Haplotype blocks can be broken between generations by recombination events, and in the absence of information from the parents, genotyping at HD full or half-siblings of the LD-target population can help identify those recombination events and increase the accuracy of imputation. Similar results have been previously reported in terrestrial species, such as in Dutch Holstein cattle (Zhang and Druet, 2010) or pigs (Cleveland and Hickey, 2013). Yoshida et al., (2018) tested a similar approach in Atlantic salmon, assessing imputation and genomic prediction accuracies for body weight in various scenarios with only siblings or parents at HD, as well as scenarios with siblings and both or only one parent in the reference population. In their study the imputation accuracy using a similar size LD-panel was significantly higher when both parents and 10% of the offspring were included in the HD-reference population, which is similar to our RPV and RP_trait_nOff scenarios that also resulted in the highest accuracy of imputation. However, in their study the scenario with only parents (similar to RPT and RPA) or only 10% of the offspring (similar to RVO) included in the HD-reference panel resulted in the lowest accuracy of imputation, showing a similar imputation accuracy for parents only and siblings only. In the current study having only the parents (RPT and RPA) resulted in a highly accurate imputation, whereas siblings only (RVO) resulted in the lowest accuracy. This difference may be explained by the lower number of parents in the HD-reference population (n=45) in Yoshida et al. (2018) compared to the current study (n=334), by the use of a previous version of the FImpute software (v2.2) or by the fact that in their study the LD-panel was constructed by selecting evenly spaced SNPs within each chromosome, while in this study the LD-panel was carefully designed for imputation (Dagnachew personal communication).

### Accuracy of genomic prediction post imputation

For all three traits in this study the accuracy of genomic prediction was higher for the HD-panel and all imputed panels than for the LD-panel or pedigree-based predictions. The genomic predictions obtained with HD, LD or imputed panels were also less biased than the pedigree-based predictions. This is in accordance with previously published literature for many different traits in various aquaculture species showing that genomic predictions obtained with GBLUP, relying on a relationship matrix built with genotype information using as low as 500 SNPs along with imputation, are always more accurate and less biased than predictions obtained with a pedigree-based BLUP and usually very close to predictions obtained with an HD-panel (Dufflocq et al., 2019; Fraslin et al., 2023; Griot et al., 2021; Kriaridou et al., 2023; Tsairidou et al., 2020; Yoshida et al., 2018, 2018). It has been reported that the magnitude of the gain in accuracy of genomic prediction comparing LD and HD panels with or without imputation may be linked with the heritability of the traits, with larger gains obtained for traits with lower heritability (Correa et al., 2017; Dufflocq et al., 2019; Ødegård et al., 2014; Tsai et al., 2017). In the current study all three traits have similar heritability (0.43 for GW, 0,46 for PD and 0.47 for CMS; see Table 2), but the gain in prediction accuracy when comparing the LD-panel and imputed scenario was higher for resistance to CMS than for the other two traits (15-21% vs 10-19% for GW and 7-11% for PD). Interestingly, for CMS the accuracy of genomic predictions obtained after imputation were not statistically different to the one obtained with the HD-panel, while for GW and resistance to PD the differences were statistically significant (see Tables S3). Since the heritabilities of these traits are similar (Table 2), the difference is likely explained by different genetic architectures. Previous studies in livestock species have reported that the inclusion in an LD-panel of SNPs with large effects or even causative variants increased the accuracy of prediction post imputation (Chen et al., 2014; da Cruz et al., 2019; Ogawa et al., 2016). In the current study growth is a polygenic trait with no significant QTL detected (Supplementary Figure S1), while resistance to PD and CMS have an oligogenic architecture with a few QTL significant at chromosome level for resistance to PD (Supplementary Figure S1; Gonen et al., 2015) and a major QTL on chromosome 27 associated with resistance to CMS (Supplementary Figure S1; Boison et al., 2019). The effect of this major QTL may not have been picked up in the LD-panel, but was probably retrieved through imputation which would explain the important gain in accuracy of genomic prediction for this trait. Indeed, on the 500 SNP array, there are 3 markers flanking the QTL region that must have allowed the correct imputation of the QTL region.

Previous studies in both terrestrial animals and fish showed the accuracy of genomic prediction post imputation was only marginally lower than the accuracy of prediction using the true HD-genotypes. However, most of those studies had high to very high (over 0.90) accuracy of imputation (Berry and Kearney, 2011; Cleveland and Hickey, 2013; Erbe et al., 2012; Tsai et al., 2017) or only simulated low errors rates of imputation from 1% to 10% (Dufflocq et al., 2019). On the other hand, some studies investigating imputation strategies resulting in less accurate imputed genotypes reported that poor imputation leads to a significant decrease in the accuracy of genomic prediction. For example, in pigs, Badke et al., (2014) showed that a less accurate imputation (R^2^=0.88) obtained using a small haplotype reference panel (n=128) resulted in a significantly lower accuracy of genomic prediction than when imputation was accurately (R^2^=0.95) performed with a large haplotype reference panel (n=1800). Grossi et al., (2018) reported a positive quadratic trend between imputation accuracy and the correlations of GEBV accuracy obtained with true (60K) or imputed genotypes. Interestingly, in Atlantic salmon Yoshida et al., (2018) compared the worst and best imputation scenarios (0.74 vs 0.94) and observed very small (3%) differences in the accuracy of genomic prediction for growth. In the current study the difference in the accuracy of imputation obtained between scenarios was not always reflected in the accuracy of genomic prediction. Indeed, when the LD-genotypes were imputed in the RP_trait_nOff scenarios, although there was an increase in the accuracy of imputation when more offspring from the different datasets were added to the HD-genotyped reference population, in general we did not see this increase reflected in the accuracy of predictions (the only exception was a small increase in prediction accuracy for GW when at least 500 individuals were added to the reference population). An even more striking example can be seen when the genotypes were imputed using the RVO scenario. Indeed, in this scenario, the accuracy of imputation was on average lower than in all other scenarios with many individuals poorly imputed (below 0.8), however for both CMS and PD the accuracy of predictions obtained after imputation were not significantly different from all other imputation scenarios. This might be explained by i) the fact that, although the quality of the average imputation was slightly degraded with this scenario, there were less individuals imputed with low accuracy; or by ii) the oligogenic architecture of the two resistance traits (Boison et al., 2019; Gonen et al., 2015); if the QTLs are correctly imputed then a lower imputation of the other markers that have little or no effect on the trait may not be important when computing the estimated breeding values (Chen et al., 2014; da Cruz et al., 2019; Ogawa et al., 2016). It is important to note that, while the GEBVs predicted after imputation with the RVO scenario are highly accurate, they are also slightly more biased than the GEBVs obtained after imputation with any other scenario. When the validation set for the prediction (i.e., selection candidates) was used alone as HD-reference population for the imputation (RVO), the estimated breeding values were slightly under-dispersed (bias >1). This under-dispersion of GEBVs could result in the identification or fewer good selection candidates.

Nevertheless, these findings are highly interesting for breeding programmes as it would suggest that the candidates for selection could be genotyped every two generations without a significant decrease in the accuracy of breeding value prediction, largely reducing the cost of genotyping without reducing the rate of genetic gain. Indeed, selection candidates from a G0 generation that would be genotyped at HD would become parents of the next generation (G1). That new generation, G1, could be genotyped only with the LD-panel and imputed to HD-level with the individuals from G0 as HD-genotyped reference population. Then the selection candidates of the next generation (G2) would need to be genotyped using an HD-panel for imputation. Thus, the cost of high-density genotyping would only be paid every two generations, i.e., every 8 years in salmon aquaculture. If the budget for genotyping is fixed, this would imply that a breeder could increase the number of selection candidates to be genotyped with an LD-panel and imputed to HD-level. The increase in the number of selection candidates would lead to an increase in selection intensity that in turn would increases the genetic gain and thus the profit of the company. The level of increase in selection intensity will depend on the price of the LD-array compared to the HD-array (and obviously the phenotypic distribution of each trait). If the LD-array is 15% cheaper than the HD-array, in theory a breeder could include 15% more selection candidates, although the additional cost of phenotyping more individuals would also need to be factored.

## Conclusion

In this Atlantic salmon population, genotype imputation from an LD-panel containing 554 carefully selected SNPs resulted in an accuracy of genomic prediction only marginally lower than that obtained with an HD-panel for three traits of interest. The makeup of the reference population used for imputation had no impact – generally, adding a proportion of the offspring to the reference population, in addition to the parents, did not result in any benefit in terms of genomic prediction accuracy. However, when one or both parents were missing in the HD-reference population, including half or full-siblings of the selection candidates in the HD-reference population increased both the accuracy of imputation and prediction. Finally, genotyping at high-density only the selection candidates and performing genotype imputation of the training population genotyped with a carefully designed LD-panel resulted in genomic prediction almost as accurate as the one obtained with all individuals genotyped at HD-level. As aquaculture breeding programmes rely on phenotypes measured on a training population composed of collaterals (full/half-sib) of the selection candidates, an imputation strategy relying on including offspring in an HD-reference panel is particularly suitable for low-cost genomic selection. This strategy could be used to only perform HD-genotyping every two generations, either reducing the cost of genotyping or increasing the selection intensity without decreasing the accuracy of prediction.

## Author contribution

**CF:** Conceptualization, Methodology, Formal analysis, Data Curation, Investigation, Writing-Original Draft, Writing-Review & Editing, Visualization. **SG:** Conceptualization, Data Curation, Writing-Review & Editing. **MB:** Conceptualization, Data Curation, Writing-Review & Editing. **SB:** Conceptualization, Data Curation, Writing-Review & Editing. **AN:** Conceptualization, Writing-Review & Editing. **DRo:** Conceptualization, Resources, Writing-Review & Editing

## Funding sources

This work is part of the AquaIMPACT project and was supported by the European Union’s Horizon 2020 research and innovation programme under the grant agreement No 818367. The study was also supported by Biotechnology and Biological Sciences Research Council (BBSRC) Institute Strategic Grants BBS/E/D/30002275, BBS/E/D/20002172, BBS/E/RL/230001A and BBS/E/RL/230001C to the Roslin Institute. DR acknowledges funding from the Axencia Galega de Innovación (GAIN, Xunta de Galicia) as part of the Oportunius programme.

## Ethics declarations

The fish used in this study are production fish from Mowi Genetics AS breeding programme. As such, the phenotypes and genotypes were collected as part of Mowi Genetics AS standard breeding practices that adhere to standards imposed by Norwegian and Irish aquaculture and fish health management authorities. Therefore, this study did not require any specific formal ethics approval.

## Competing interest

AN, SB, MB and SG are currently or were employed by Mowi Genetics AS at the time of the study. CF and DR declare that they have no competing interests.

## Data availability

The data utilized in this study were provided by Mowi Genetics AS and are not publicly accessible.

## Supplementary data

**Table S1. P-values of suggestive or not-significant pairwise Dunn tests comparing the accuracy of imputation obtained for all scenarios, corrected for multiple testing with Benjamini-Hochberg**.

*Suggestive comparison when corrected 0.05 < p_value < 0.01.

*** Significant comparison with a corrected p-value < 0.001.

**RPT**: Reference Parents Trait. Imputation was performed with the reference population composed of all parents with imputation performed on offspring per trait.

**RPA**: Reference Parents All. Imputation was performed with the reference population composed of all parents, all offspring from the three traits imputed together.

**RPV**: Reference Parents Validation. Imputation was performed with the reference population composed of all parents and the offspring from the validation set (1/10^th^ of the offspring) per trait.

**RVO**: Reference Validation Only. Imputation was performed with the reference population composed of the offspring from the validation set (1/10^th^ of the offspring) per trait.

**RP_trait_n:** Reference Parents trait. Imputation was performed with the reference population composed of all parents and n (100, 300, 700 or 1000) offspring randomly sampled from the growth dataset.

**GW**: Growth, **PD**: Pancreatic Disease, **CMS**: Cardiomyopathy syndrome.

**Table S2. Number and proportion (%) of individual with an average imputation accuracy below 0.8 and below 0.5**

RPT: Reference Parents Trait. Imputation was performed with the reference population composed of all parents with imputation performed on offspring per trait.

RPA: Reference Parents All. Imputation was performed with the reference population composed of all parents, all offspring from the three traits imputed together.

RPV: Reference Parents Validation. Imputation was performed with the reference population composed of all parents and the offspring from the validation set (1/10^th^ of the offspring) per trait.

RVO: Reference Validation Only. Imputation was performed with the reference population composed of the offspring from the validation set (1/10^th^ of the offspring) per trait.

RP_trait_nOff: Reference Parents trait. Imputation was performed with the reference population composed of all parents and n (100, 300, 700 or 1000) offspring randomly sampled from the growth dataset.

GW: Growth, PD: Pancreatic Disease, CMS: Cardiomyopathy syndrome.

**Table S3. P-values of pairwise Dunn tests comparing the accuracy of genomic prediction obtained for all scenarios, corrected for multiple testing with Benjamini-Hochberg**.

HD: Prediction obtained with the high-density panel (47K SNPs).

LD: Prediction obtained with the un-imputed low-density panel (554 SNPs).

PED: Prediction obtained with a pedigree-based approach.

RPT: Reference Parents Trait. Imputation was performed with the reference population composed of all parents with imputation performed on offspring per trait.

RPA: Reference Parents All. Imputation was performed with the reference population composed of all parents, all offspring from the three traits imputed together.

RPV: Reference Parents Validation. Imputation was performed with the reference population composed of all parents and the offspring from the validation set (1/10^th^ of the offspring) per trait.

RVO: Reference Validation Only. Imputation was performed with the reference population composed of the offspring from the validation set (1/10^th^ of the offspring) per trait.

GW: Growth, PD: Pancreatic Disease, CMS: Cardiomyopathy syndrome.

**Table S4. P-values of pairwise Dunn tests comparing the accuracy of genomic predictions obtained for all RP_trait_nOff scenarios, corrected for multiple testing with Benjamini-Hochberg**.

Table S4a for GW, Table S4b for PD and Table S4c for CMS

GW: Growth, PD: Pancreatic Disease, CMS: Cardiomyopathy syndrome.

**Supplementary Figures S1. Manhattan plots representing the GWAS outputs for each trait, growth (GW; in blue), resistance to pancreatic disease (PD; in red) and to cardiomyopathy syndrome (CMS; in green)**. GWAS were performed with the GCTA software using the mixed linear BLUP-animal model described in the methods, with sex as fixed effect in the model for GW.

The red line is the 5% Bonferroni correction threshold at the genome wide level: -log_10_(0.05/47,061). The blue line is the 5% Bonferroni correction threshold at the chromosome wide level: -log_10_(0.05/(47,061/29)).

## Notes

### Competing Interest Statement

Ashie Norris , Solomon Boison, Matthew Baranski and Serap Gonen were employed by Mowi Genetics AS at the time of the study.
Ashie Norris , Solomon Boison and Matthew Baranski are employed by Mowi Genetics AS.

