## Supplementary Figure 1 for "Optimization of reference population for imputation of low-density SNPs panel for genomic prediction in Atlantic salmon"

**Supplementary Figures S1. Manhattan plots representing the GWAS outputs for each trait, growth (GW; in blue), resistance to pancreatic disease (PD; in red) and to cardiomyopathy syndrome (CMS; in green).** GWAS were performed with the GCTA software using the mixed linear BLUP- animal model described in the methods, with sex as fixed effect in the model for GW.

The red line is the 5% Bonferroni correction threshold at the genome wide level: -log_10_(0.05/47,061). The blue line is the 5% Bonferroni correction threshold at the chromosome wide level: -log_10_(0.05/(47,061/29)).
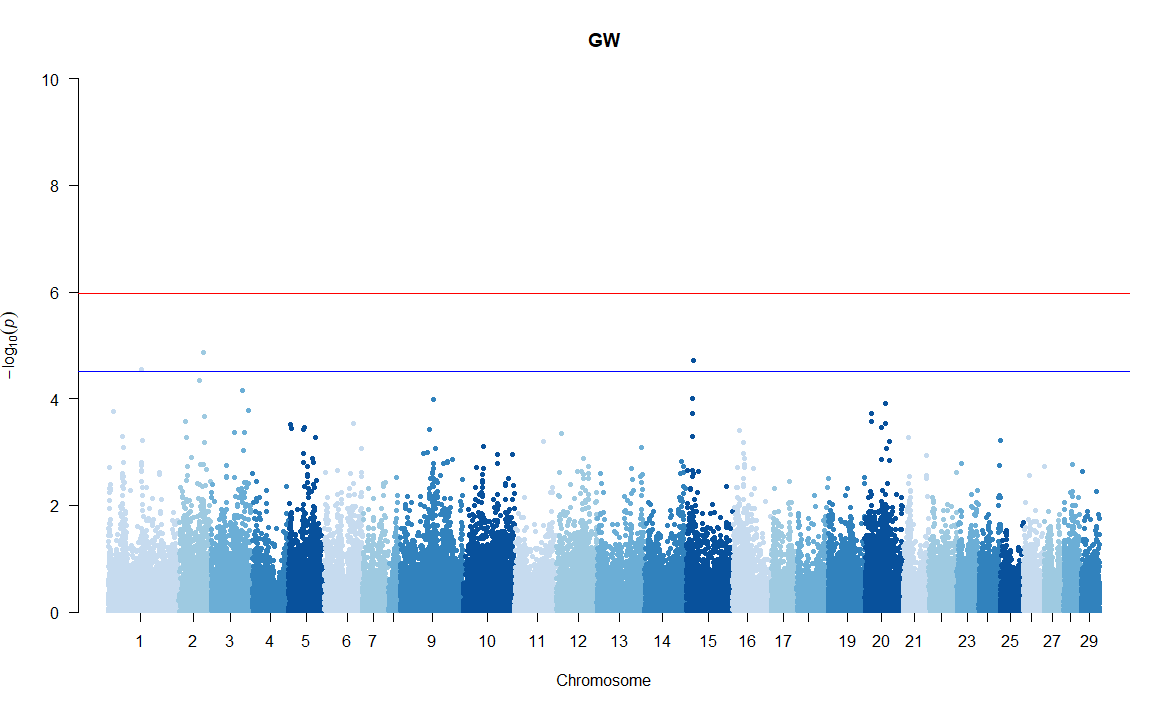

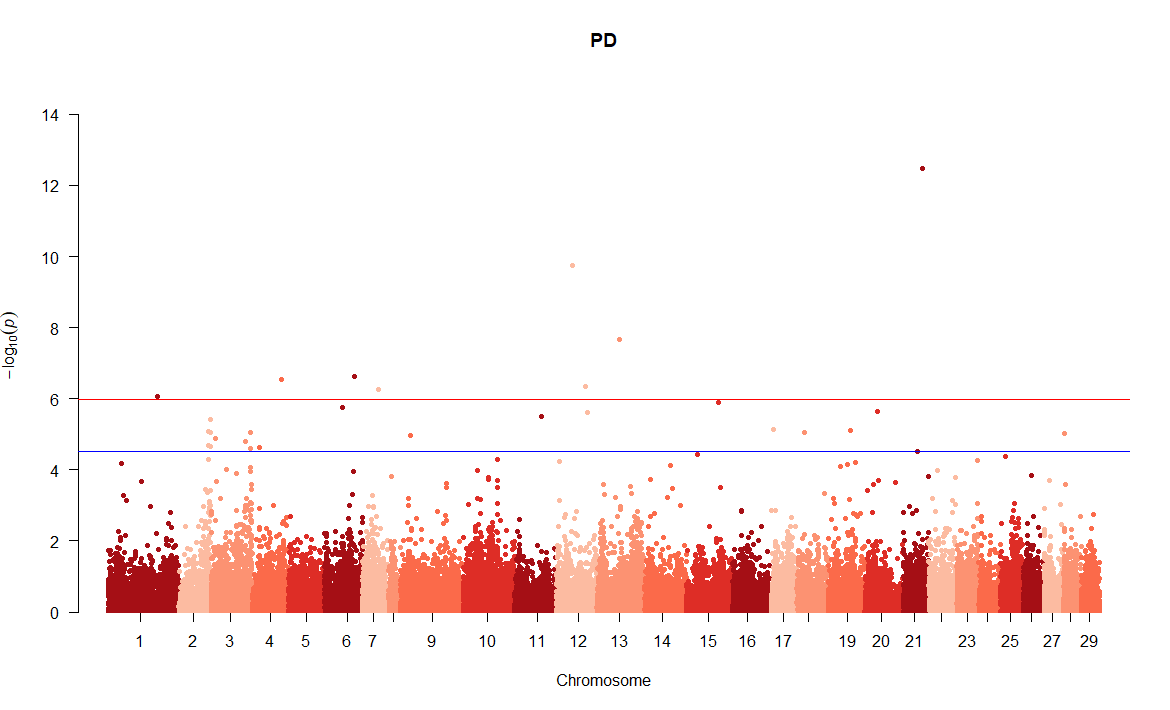

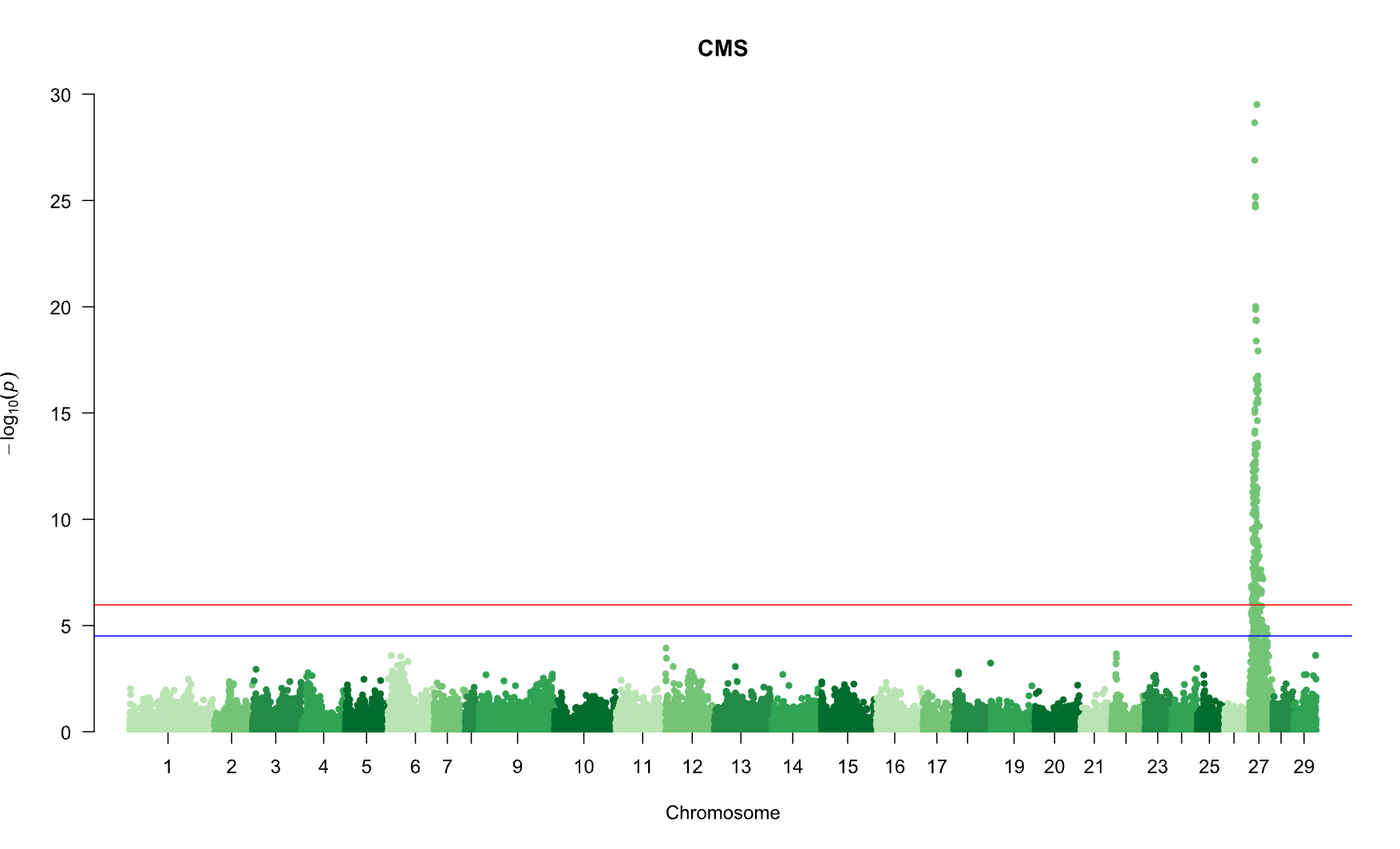
