## Supplementary Tables 1, 2, 3 for "Optimization of reference population for imputation of low-density SNPs panel for genomic prediction in Atlantic salmon"

**Table S1. P-values of suggestive or not-significant pairwise Dunn tests comparing the accuracy of imputation obtained for all scenarios, corrected for multiple testing with Benjamini-Hochberg.**

| **Trait imputed** | **Imputation scenarios** | | **P-value** |
| --- | --- | --- | --- |
| **GW** | RVO | RPV | 0*** |
| **GW** | RVO | RPA | 3.08 e-303*** |
| **GW** | RVO | RPT | 0*** |
| **GW** | RPV | RPA | 0*** |
| **GW** | RPV | RPT | 1.4 e-306*** |
| **GW** | RPA | RPA | 1.1 e-05*** |
| **PD** | RVO | RPV | 0*** |
| **PD** | RVO | RPA | 1.01 e-140*** |
| **PD** | RVO | RPT | 9.29 e-195*** |
| **PD** | RPV | RPA | 1.87 e-170*** |
| **PD** | RPV | RPT | 3.49 e-120*** |
| **PD** | RPA | RPA | 2.79 e-06*** |
| **CMS** | RVO | RPV | 0*** |
| **CMS** | RVO | RPA | 3.2 e-132*** |
| **CMS** | RVO | RPT | 2.906 e-168*** |
| **CMS** | RPV | RPA | 4.86 e-126*** |
| **CMS** | RPV | RPT | 3.58 e-95*** |
| **CMS** | RPA | RPT | 5.7 e-04*** |
| **GW** | RP_CMS_700 | RP_CMS_500 | 0.267 |
| **GW** | RP_CMS_700 | RP_PD_700 | 0.255 |
| **PD** | RP_CMS_500 | RP_GW_500 | 0.031* |
| **PD** | RP_CMS_700 | RP_GW_700 | 0.198 |
| **PD** | RP_CMS_100 | RP_PD_100 | 0.012* |
| **PD** | INP_GW_100 | RP_PD_100 | 0.374 |
| **PD** | RP_CMS_700 | RP_PD_500 | 0.463 |
| **PD** | RP_GW_700 | RP_PD_500 | 0.227 |
| **CMS** | RP_CMS_100 | RP_GW_100 | 0.049* |
| **CMS** | RP_CMS_300 | RP_GW_300 | 0.277 |
| **CMS** | RP_CMS_500 | RP_GW_700 | 0.368 |
| **CMS** | RP_CMS_700 | RP_GW_1000 | 0.032* |
| **CMS** | RP_CMS_100 | RP_PD_100 | 0.408 |
| **CMS** | RP_CMS_300 | RP_PD_300 | 0.400 |
| **CMS** | RP_CMS_500 | RP_PD_700 | 0.471 |
| **CMS** | RP_GW_100 | RP_PD_100 | 0.079 |
| **CMS** | RP_GW_300 | RP_PD_300 | 0.373 |
| **CMS** | RP_GW_500 | RP_PD_500 | 0.377 |
| **CMS** | RP_GW_700 | RP_PD_700 | 0.349 |

*Suggestive comparison when corrected 0.05 < p_value < 0.01.

*** Significant comparison with a corrected p-value < 0.001.

**RPT**: Reference Parents Trait. Imputation was performed with the reference population composed of all parents with imputation performed on offspring per trait.

**RPA**: Reference Parents All. Imputation was performed with the reference population composed of all parents, all offspring from the three traits imputed together.

**RPV**: Reference Parents Validation. Imputation was performed with the reference population composed of all parents and the offspring from the validation set (1/10^th^ of the offspring) per trait.

**RVO**: Reference Validation Only. Imputation was performed with the reference population composed of the offspring from the validation set (1/10^th^ of the offspring) per trait.

**RP_trait_n:** Reference Parents trait. Imputation was performed with the reference population composed of all parents and n (100, 300, 700 or 1000) offspring randomly sampled from the growth dataset.

**GW**: Growth, **PD**: Pancreatic Disease, **CMS**: Cardiomyopathy syndrome

**Table S2. Number and proportion (%) of individual with an average imputation accuracy below 0.8 and below 0.5**

|  | GW | | PD | | CMS | |
| --- | --- | --- | --- | --- | --- | --- |
|  | **0.5** | **0.8** | **0.5** | **0.8** | **0.5** | **0.8** |
| RPT | 10  (0.33%) | 350  (11.7%) | 7 (0.48%) | 188  (13.0%) | 6  (0.55%) | 104  (9.5%) |
| RPA | 6  (0.20%) | 351  (11.7%) | 2 (0.14%) | 189  (13.1%) | 2  (0.18%) | 104  (9.5%) |
| RPV | 7  (0.26%) | 254  (9.4%) | 4 (0.31%) | 182  (14.0%) | 3  (0.32%) | 85  (9.2%) |
| RVO | 1  (0.04%) | 1351  (50.2%) | 0  (0%) | 1223  (93.9%) | 0  (0%) | 816  (88.5%) |
| RP_GW  100 | 8  (0.28%) | 347  (13.4%) | 7 (0.48%) | 187  (12.9%) | 4  (0.37%) | 103  (9.4%) |
| RP_GW  300 | 7  (0.26%) | 243  (10.2%) | 4 (0.28%) | 141  (9.7%) | 2  (0.18%) | 81  (7.4%) |
| RP_GW  500 | 8  (0.32%) | 203  (9.3%) | 4 (0.28%) | 118  (8.1%) | 2  (0.18%) | 69  (6.3%) |
| RP_GW  700 | 7  (0.30%) | 160  (8.0%) | 3 (0.21%) | 96  (6.6%) | 1  (0.09%) | 57  (5.2%) |
| RP_GW  1000 | 7  (0.35%) | 112  (6.63%) | 3 (0.21%) | 78  (5.4%) | 1  (0.09%) | 47  (4.3%) |
| RP_PD  100 | 4  (0.15%) | 347  (12.9%) | 4  (0.28%) | 186  (12.85%) | 2  (0.18%) | 103  (9.41%) |
| RP_PD  300 | 5  (0.19%) | 225  (8.36%) | 4  (0.28%) | 130  (8.98%) | 1  (0.09%) | 74  (6.76%) |
| RP_PD  500 | 6  (0.22%) | 176  (6.54%) | 4  (0.28%) | 92  (6.35%) | 1  (0.09%) | 54  (4.94%) |
| RP_PD  700 | 7  (0.26%) | 143  (5.32%) | 4  (0.28%) | 76  (5.25%) | 2  (0.18%) | 46  (4.20%) |
| RP_CMS  100 | 6  (0.22%) | 347  (12.90%) | 5  (0.35%) | 187  (12.91%) | 4  (0.37%) | 103  (9.41%) |
| RP_CMS  300 | 4  (0.15%) | 277  (10.30%) | 3  (0.21%) | 152  (10.50%) | 2  (0.18%) | 87  (7.95%) |
| RP_CMS  500 | 4  (0.15%) | 236  (8.77%) | 3  (0.21%) | 115  (7.94%) | 0  (0%) | 71  (6.49%) |
| RP_CMS  700 | 6  (0.22%) | 190  (7.06%) | 3  (0.21%) | 95  (6.56%) | 0  (0%) | 58  (5.30%) |

**RPT**: Reference Parents Trait. Imputation was performed with the reference population composed of all parents with imputation performed on offspring per trait.

**RPA**: Reference Parents All. Imputation was performed with the reference population composed of all parents, all offspring from the three traits imputed together.

**RPV**: Reference Parents Validation. Imputation was performed with the reference population composed of all parents and the offspring from the validation set (1/10^th^ of the offspring) per trait.

**RVO**: Reference Validation Only. Imputation was performed with the reference population composed of the offspring from the validation set (1/10^th^ of the offspring) per trait.

**RP_trait_n**: Reference Parents trait. Imputation was performed with the reference population composed of all parents and n (100, 300, 700 or 1000) offspring randomly sampled from the growth dataset.

**GW**: Growth measured as gutted weight. **PD**: Pancreas disease. **CMS**: Cardiomyopathy syndrome.

**Table S3. P-values of Kruskal-Wallis rank tests comparing the accuracy of genomic prediction obtained for all scenarios, corrected for multiple testing with Benjamini-Hochberg.**

| **GW** | **HD** | **RPA** | **RPT** | **RPV** | **RVO** | **LD** |
| --- | --- | --- | --- | --- | --- | --- |
| RPA | 1.621252e-03 |  |  |  |  |  |
| RPT | 3.030054e-03 | 4.077895e-01 |  |  |  |  |
| RPV | 5.228620e-02 | 9.799215e-02 | 1.388619e-01 |  |  |  |
| RVO | 1.536341e-09 | 1.760254e-03 | 9.116320e-04 | 1.185587e-05 |  |  |
| LD | 1.669951e-25 | 9.986309e-14 | 1.862176e-14 | 2.180842e-18 | 7.844027e-06 |  |
| PED | 1.108576e-43 | 1.234449e-27 | 1.211415e-28 | 3.265241e-34 | 2.207676e-15 | 3.487503e-04 |
| **PD** | **HD** | **RPA** | **RPT** | **RPV** | **RVO** | **LD** |
| RPA | 2.426712e-04 |  |  |  |  |  |
| RPT | 3.120645e-04 | 4.655860e-01 |  |  |  |  |
| RPV | 1.029008e-04 | 4.267541e-01 | 4.126047e-01 |  |  |  |
| RVO | 9.636309e-07 | 1.108445e-01 | 1.004502e-01 | 1.568379e-01 |  |  |
| LD | 1.150658e-14 | 2.043714e-05 | 1.552251e-05 | 5.204590e-05 | 2.210226e-03 |  |
| PED | 4.323992e-28 | 1.029780e-13 | 7.040129e-14 | 4.950306e-13 | 8.638814e-10 | 8.090783e-04 |
| **CMS** | **HD** | **RPA** | **RPT** | **RPV** | **RVO** | **LD** |
| RPA | 4.503272e-01 |  |  |  |  |  |
| RPT | 1.654596e-01 | 1.899625e-01 |  |  |  |  |
| RPV | 4.508354e-01 | 4.211199e-01 | 1.500496e-01 |  |  |  |
| RVO | 1.675306e-01 | 1.448157e-01 | 1.771051e-02 | 1.854710e-01 |  |  |
| LD | 1.719197e-10 | 8.581179e-11 | 6.470586e-14 | 4.849195e-10 | 1.456928e-07 |  |
| PED | 3.717239e-33 | 1.209347e-33 | 5.082078e-39 | 2.398975e-32 | 1.881908e-27 | 1.468906e-08 |

**HD**: Prediction obtained with the high-density panel (47K SNPs).

**LD**: Prediction obtained with the un-imputed low-density panel (554 SNPs).

**PED**: Prediction obtained with a pedigree-based approach.

**RPT**: Reference Parents Trait. Imputation was performed with the reference population composed of all parents with imputation performed on offspring per trait.

**RPA**: Reference Parents All. Imputation was performed with the reference population composed of all parents, all offspring from the three traits imputed together.

**RPV**: Reference Parents Validation. Imputation was performed with the reference population composed of all parents and the offspring from the validation set (1/10^th^ of the offspring) per trait.

**RVO**: Reference Validation Only. Imputation was performed with the reference population composed of the offspring from the validation set (1/10^th^ of the offspring) per trait.

**GW**: Growth, **PD:** Pancreatic Disease, **CMS**: Cardiomyopathy syndrome.
